# Molecular mechanisms underlying attenuation of live attenuated Japanese encephalitis virus vaccine SA14-14-2

**DOI:** 10.1101/2021.01.14.426643

**Authors:** Pooja Hoovina Venkatesh, Saurabh Kumar, Naveen Kumar, Krishna Chaitanya, Lance Turtle, Vijaya Satchidanandam

## Abstract

The live attenuated Japanese encephalitis virus vaccine SA14-14-2 demonstrated ≥ 95 % efficacy and is today the vaccine of choice against JEV globally. Relative to its parent strain SA14, SA14-14-2 carries 46 nucleotide and 24 amino acid alterations, with 8 of the latter located within the envelope glycoprotein. The vaccine strain also fails to synthesize the nonstructural protein NS1’ owing to a silent mutation that abrogates a-1-frameshifting event close to the 5’ end of the NS2A coding sequence. Previous studies employing reverse genetics and mouse models implicated both absence of NS1’ and mutated E, in attenuation of SA14-14-2. We demonstrate progressive reduction in ER stress sensor PERK levels and increased expression of CEBP-homologous protein (CHOP), accompanied by dephosphorylation of eIF2α, inhibition of autophagy maturation and necroptosis following infection of cultured cells with wild-type JEV strain P20778. Autonomous expression of NS1’ caused constitutive up-regulation of CHOP and loss of PERK. Conversely, infection with SA14-14-2 led to significantly increased IRE-1α activation, ER chaperone levels and autophagy. We report labile conformational epitopes accompanied by drastically reduced folding kinetics of intracellular SA14-14-2 envelope protein engendered by sluggish oxidation of cysteine sulfhydryl groups to form disulfide bonds within the endoplasmic reticulum along with altered envelope epitopes in extracellular SA14-14-2 viral particles. We also demonstrate near total conversion of prM to pr and M in SA14-14-2 virus particles. These alterations were accompanied by enhanced activation of mouse and human antigen presenting cells by SA14-14-2 along with superior CD8^+^ recall T cell responses to viral structural proteins in volunteers vaccinated with SA14-14-2.

**Author Summary:** The random process of cell culture passage adopted in generation of most live attenuated virus vaccines leads to fixation of multiple nucleotide changes in their genomes and renders it difficult if not impossible to pinpoint those mutations primarily responsible for their attenuated phenotype. Identifying the precise attenuating mutations and their *modi operandi* should aid in developing rationally attenuated vaccines for other viruses. We discovered that wild type (WT) JEV uses the nonstructural protein NS1’ to take over the host protein synthesis machinery to produce viral proteins. Loss of NS1’ in SA14-14-2 deprives the vaccine strain of this ability. Viruses uniformly target host death pathways to avoid generating potent antiviral immune responses. WT JEV prevents autophagy maturation. Conversely the SA14-14-2 vaccine activates autophagy due to unresolved ER stress caused by inability of its envelope glycoprotein to fold promptly post synthesis. Combined with enhanced proteolytic cleavage of the viral prM protein in SA14-14-2, this resulted in altered envelope epitopes on extracellular SA14-14-2 virus particles. These changes culminated in enhanced activation of innate and adaptive immune responses by SA14-14-2.

## INTRODUCTION

The genus *Flavivirus,* in the family Flaviviridae, comprising numerous vector-borne human pathogens has spawned two highly efficacious live attenuated vaccines in Yellow Fever virus (YFV)-17D and Japanese encephalitis virus (JEV)-SA14-14-2, both of which have contributed to significant reduction of disease incidence from their respective viral pathogens. Flavivirus genomes are single strand RNA of positive polarity and encode a single large polyprotein which is processed by host and virally encoded proteases to give rise to 3 structural (capsid, C; envelope, E; and premembrane, prM) and 7 non-structural (NS; 1, 2A, 2B, 3, 4A, 4B and 5) proteins. JEV is represented by a single serotype and 5 genotypes [1] and was first isolated in Japan in 1934, giving us the prototype Nakayama strain [2]. Based on meta-analysis of published literature and national incidence estimates of 124 countries the annual global incidence of JEV encephalitis was computed around 67,900 [3].

Of the 24 amino acid changes identified in SA14-14-2 relative to its parent Chinese strain SA14, the largest number, namely 8 were located within the envelope glycoprotein E [4, 5]. Similarly, 8 of the 22 amino acid changes in YF17D relative to its parent Asibi strain were found in the E protein [6]. Studies in the mouse model implicated multiple mutations within the E gene of SA14-14-2 in the attenuated phenotype of this vaccine strain [7–10]. Additionally, a silent G to A mutation near the beginning of the NS2A gene which led to abrogation of-1 ribosome frameshifted NS1’ synthesis was reported to determine loss of neurovirulence of SA14-14-2 [11]. Both viral E and NS1’ are secreted glycoproteins synthesized on endoplasmic reticulum (ER)-bound ribosomes, that are dependent on the ER lumen for glycosylation and folding. The molecular mechanisms underlying the attenuation caused by these mutations have not been elucidated.

As obligate intracellular parasites, viruses rely on usurping the host translational machinery for their survival and propagation. The dominant immediate host response to viral infection is a transient shutdown of translation, which serves to limit viral protein synthesis [12]. PKR-like ER resident kinase (PERK), one of the three ER resident stress sensors which attenuates translation via phosphorylation of the downstream eukaryotic translation initiation factor eIF2α constitutes one of the earliest pathways that responds to the increased burden of viral protein synthesis within the ER. PERK activation was reported to have divergent effects on replication of viruses; whereas activated PERK negatively regulated Transmissible Gastroenteritis Virus (TGEV) replication [13], translation of dengue proteins in mosquito cells continued despite PERK activation which in fact prolonged survival of infected cells, ultimately aiding viral replication [14]. Inhibition/degradation of its cytoplasmic counterpart PKR by multiple viruses to aid their survival and replication has been extensively documented [15–21]. Sustained phosphorylation of eIF2α and the resultant translation block is known to activate autophagy [22]. Several viruses have evolved mechanisms both to prevent phosphorylation of eIF2α and leverage autophagy for their own benefit. Influenza A virus, coxsackie B virus and rotavirus have all evolved to increase autophagy flux and utilize autophagosomal membranes to facilitate their replication while simultaneously preventing the antiviral effects of late stages of autophagy maturation and lysozome fusion of autophagosomes [reviewed in [23]]. Thus, viruses escape completion of autophagy and its attendant activation of inflammation leading to efficient priming of cytolytic CD8^+^ T cells. Autophagy has been demonstrated to facilitate efficient transporter of antigenic peptide (TAP)-independent presentation of viral antigenic peptides contained within autophagolysosomes by fusion with endosomes containing internalized MHC Class I molecules [24].

The most conserved ER stress sensor IRE-1α, an ER transmembrane kinase and endoribonuclease which is activated when unfolded proteins accumulate within the ER, serves to match protein folding burden within the ER with capacity. IRE-1α generates the transcription factor XBP-1 [25] through nonconventional cytoplasmic splicing which in turn elps to maintain ER homeostasis and prevent activation of cell death pathways caused by sustained ER stress. The IRE-1α-XBP-1 axis was reported to be vital and essential for development and survival of dendritic cells [26], certain subsets of which were shown to constitutively activate the IRE-1α-XBP-1 pathway [26, 27]. More importantly, both transcription and splicing of XBP-1 mRNA were reported to be increased in response to viral or bacterial infection within antigen specific CD8^+^ T cells, whose differentiation into killer cell lectin-like receptor G1 (KLRG1)-expressing terminal effector cells was dependent on XBP-1 expression [28]. Thus, IRE-1α is poised to link infection-induced ER perturbation and unfolded protein response (UPR) to subsequent steps of antigen presentation and generation of host-protective immune responses.

In order to understand the molecular mechanisms by which mutations in E and NS1’ glycoproteins of JEV contribute to the attenuation and immunogenic efficacy of SA14-14-2, we compared markers of ER stress, UPR and death pathways in cells infected with the WT (P20778) and vaccine (SA14-14-2) strains of JEV. Our results highlighted the role of JEV NS1’ in orchestrating dephosphorylation of phosphor e-IF2α by modulating both upstream kinase PERK and feedback dephosphorylation by CAAT/enhancer binding protein (CEBP) homologous protein (CHOP) to achieve efficient viral protein synthesis. Sluggish folding kinetics of SA14-14-2 E protein and the resultant aggravated and unresolved ER protein folding strain led to enhanced autophagy in SA14-14-2 infected cells. The consequent efficient cross presentation of viral antigen-containing autophagic vesicles most likely resulted in the observed augmented activation of antigen presenting cells by SA14-14-2 and enhanced recall CD8^+^ T cell responses to viral structural proteins envelope and capsid in individuals vaccinated with SA14-14-2.

## RESULTS

### Differential modulation of PERK-eIF2α-CHOP pathway by wild type and vaccine strains of JEV

As a first step to query the underlying basis for attenuation of SA14-14-2 we compared activation of the ER stress sensor PERK in JEV-infected cells. We observed progressive reduction in levels of total and phosphorylated PERK in Neuro2A (N2A; Fig 1A, S1Fig D) as well as porcine kidney fibroblasts (PS; S1 Fig A) along with rapid dephosphorylation of eIF2α in both cell lines following infection with P20778 (Fig 1B, S1 Fig E, B and C). Infection with SA14-14-2 in contrast, led to sustained increase in phosphorylation of eIF2α in both cell lines (Fig 1B, S1 Fig B, C and E). We also observed impressive upregulation of CHOP which mediates induction of the regulatory subunit of the phosphatase responsible for eIF2α dephosphorylation, accompanied by the transcription factor ATF4, only in cells infected with P20778 (Fig 1C and D, S1Fig F). CHOP and ATF4 upregulation in the absence of eIF2α phosphorylation, required to override the upstream ORF in the CHOP and ATF4 mRNA [29] was intriguing; we did not investigate a potential role for the reported phosphorylation of eIF4E [30] in CHOP expression. Thus, wild type JEV efficiently prevented the phosphor-eIF2α-mediated translation block triggered by the host upon viral infection by destroying upstream PERK and stimulating downstream feedback expression of CHOP to facilitate viral protein synthesis. Interestingly, the ER stress sensor IRE-1α revealed phosphorylation-mediated activation only in SA14-14-2-infected cells, accompanied by enhanced expression of XBP-1 (Fig 1E, S1 Fig G and H), pointing to increased protein folding stress and activation of UPR following infection with SA14-14-2 but not P20778.

**Figure 1.**
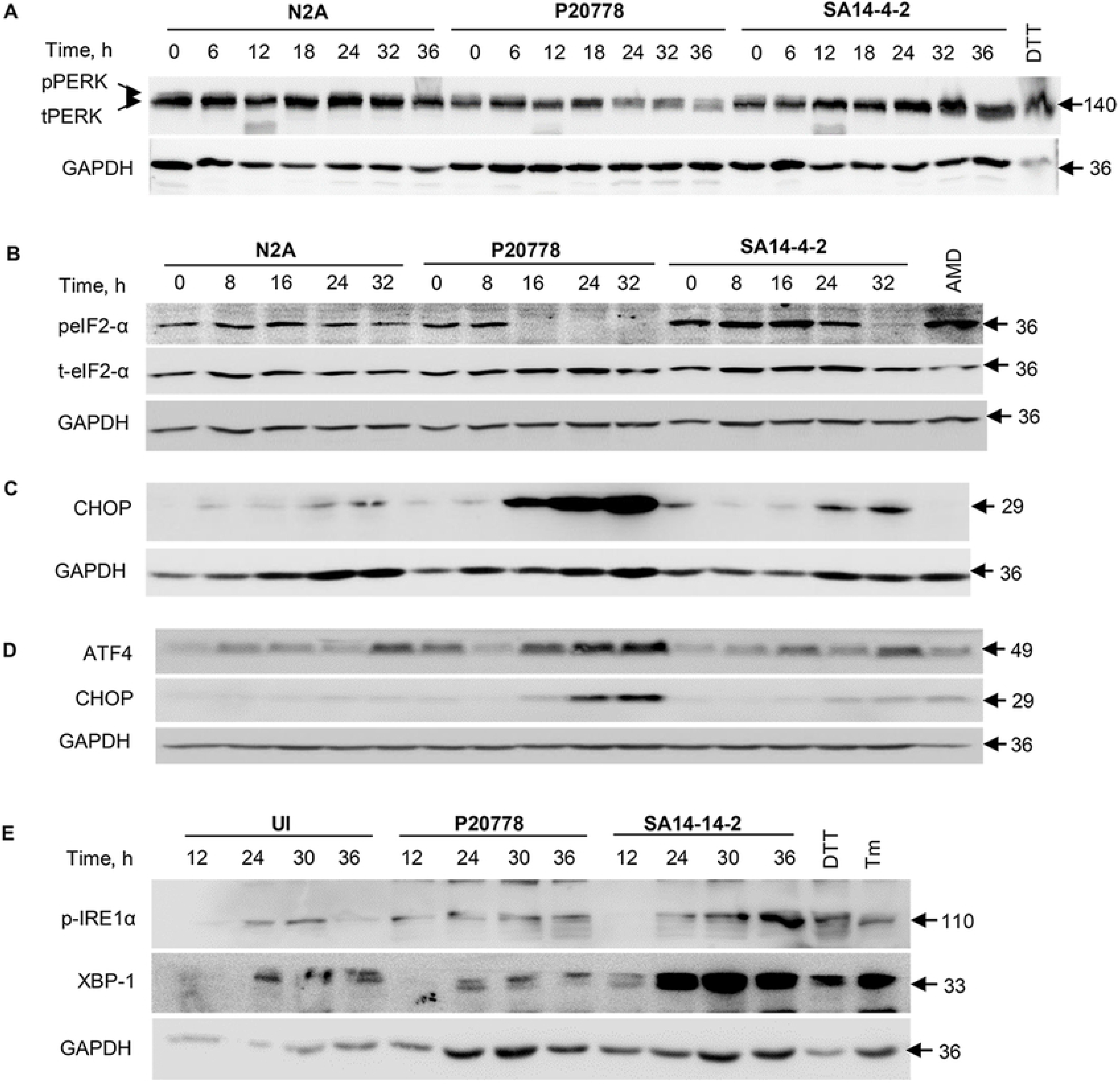
Differential modulation of the PERK-eIF2α pathway by WT and vaccine strains of JEV. (A) Progressive loss of PERK in WT JEV-infected cells. N2A cells were infected with P20778 and SA14-14-2 at a multiplicity of 10 and lysate equivalent to 2 × 10^5^ cells electrophoresed on SDS-10 % PAGE and transferred onto nitrocellulose membrane were immunoblotted with antibodies specific to PERK and GAPDH. 10 mM DTT served as a positive control. DTT-3 h treatment of cells with 10 mM DTT. Both phosphorylated and non-phosphorylated forms of PERK detected by the antibody are indicated. (B) Sequential immunoblotting of virus infected cell lysates harvested at the time points indicated, with antibodies specific to peIF2α, total eIF2α and GAPDH. AMD-cells treated for 18 h with 5 μg/ml AMD. Immunoblotting of virus infected cell lysates harvested at the time points indicated with antibodies specific to (C) CHOP and GAPDH, (D) ATF4, CHOP and GAPDH and (E) p-IRE1α, XBP-1 and GAPDH. DTT-3 h treatment of cells with 10 mM DTT; Tm-5 h treatment of cells with 5 μg/ml tunicamycin. Numbers on the right indicate sizes in kilo Daltons (kDa), of proteins detected in each panel.

### JEV NS1’ activates CHOP expression

To query a role for NS1’ (absent in SA14-14-2) in modulating the PERK pathway, we created cells stably expressing NS1 or NS1’ of P20778 by lentivirus transduction. PS cells stably expressing NS1’ revealed constitutive high expression of CHOP (Fig 2, lanes 9 to 12). Cells expressing P20778 NS1 protein were distinctly devoid of CHOP, even on long exposure (Fig 2, lanes 5 to 8). Surprisingly, NS1’-induced CHOP expression did not lead to dephosphorylation of eIF2α, presumably due to vital compensatory pathways in cells conditioned to constitutively express CHOP over multiple passages. We also observed progressive loss of PERK levels in NS1’-expressing cells over a span of 36 hours (Fig 2, lanes 9 to 12). These results revealed a role for the C-terminal frameshifted 52 amino acid segment of NS1’ in modulating levels of CHOP and PERK following viral infection. Loss of NS1’ in SA14-14-2-infected cells would understandably cripple the vaccine strain due to its inability to appropriate the translation machinery of the host.

**Figure 2.**
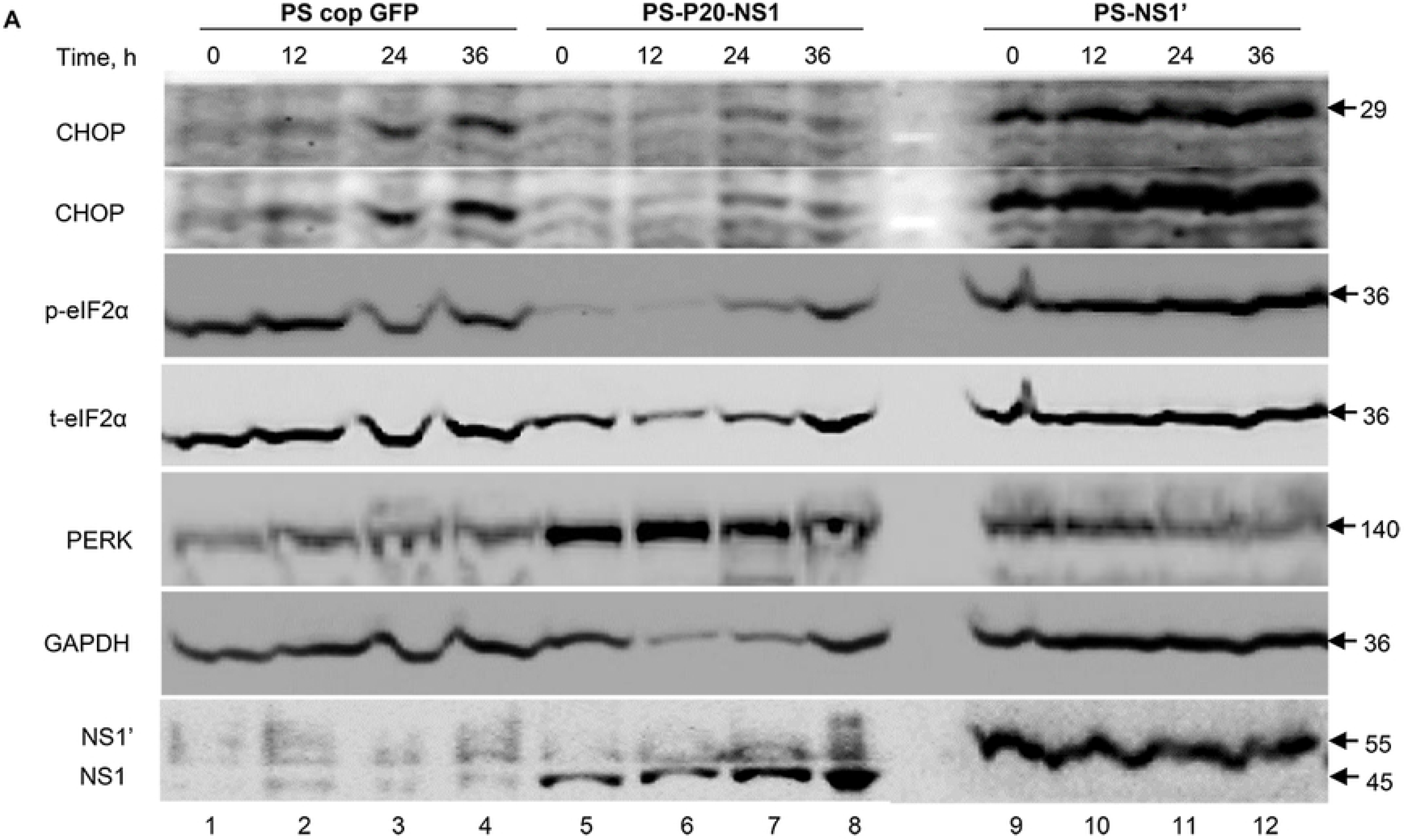
JEV NS1’ protein induces constitutive expression of CHOP. PS cells transduced with lentiviruses stably expressing codon optimized GFP (PS-cop-GFP), JEV P20778 NS1 protein (PS-P20-NS1) or JEV P20778 NS1’ protein (PS-P20-NS1’) were harvested at indicated time points and immunoblotted with antibodies to the proteins indicated on the left. JEV proteins NS1 and NS1’ were detected using the mAb H5D12 and CB1A2, respectively generated in this study. Longer exposure did not reveal CHOP expression in NS1-expressing cells. Numbers on the right indicate sizes in kDa, of proteins detected in each panel.

### WT JEV blocks autophagy maturation

The sustained high phosphorylation levels of eIF2α in SA14-14-2-infected cells led us to investigate autophagy. Diverse reports exist, of the positive as well as negative effect of autophagy on JEV replication [31, 32]. P20778 induced impressive stabilization and consequent increased levels of lipidated LC3-II and p62 beyond 16 h of infection, not seen for SA14-14-2 (Fig 3A, S2 Fig C and D). In contrast we observed progressive loss of Beclin-1, a crucial protein required to generate the autophagy maturation complex in P20778-, but not SA14-14-2-infected PS cells (S2 FigA). Thus, wild type JEV appears to block autophagy maturation by targeting Beclin in a manner reminiscent of influenza A virus whose M2 protein directly interacts with Beclin to disrupt the autophagy maturation complex [33]. We then queried any changes in autophagy flux in virus infected cells using the inhibitor bafilomycin. Addition of bafilomycin stabilized and increased LC3-II levels in P20778-infected cells at early times post infection, revealing a modest increase in autophagy flux at 12 and 24 h post infection (Fig 3B, compare lanes 3 and 4 with 13 and 14, S2 FigE). A role for autophagy early during JEV infection was also reported previously [31]. However, we observed no further stabilization in levels of LC3-II by bafilomycin over that already achieved by WT-JEV beyond 24 h post infection (Fig 3B, compare lanes 5 and 6 with 15 and 16, S2 FigE). In contrast, bafilomycin revealed impressive and sustained increase in autophagy flux within cells infected with SA14-14-2 (Fig 3B, compare lanes 8 to 10 with 18 to 20, S2 FigE), corroborated by increased levels of ATG5 and ATG7 (Fig 3C). Thus, WT JEV while increasing autophagy during early stages of replication, clearly prevented autophagy maturation and late-stage degradative loss of LC3-II and p62. Such a strategy would provide JEV with the ER-derived membranes that host the viral replicase complex [34–36] while at the same time preventing the autophagy-induced efficient priming of virus-specific CD8^+^ T cells [23]. This ability to commandeer the host autophagy pathway was obviously lost during attenuation of SA14-14-2; activated autophagy flux in SA14-14-2-infected cells is perhaps triggered by persistently phosphorylated eIF2α [22]. We did not observe alterations in levels of LC3-II or p62 in cells stably expressing NS1’ (data not shown). The viral protein(s) responsible for stabilization of LC3-II and p62 and the mutation(s) in SA14-14-2 responsible for abrogating this effect are yet to be determined.

**Figure 3.**
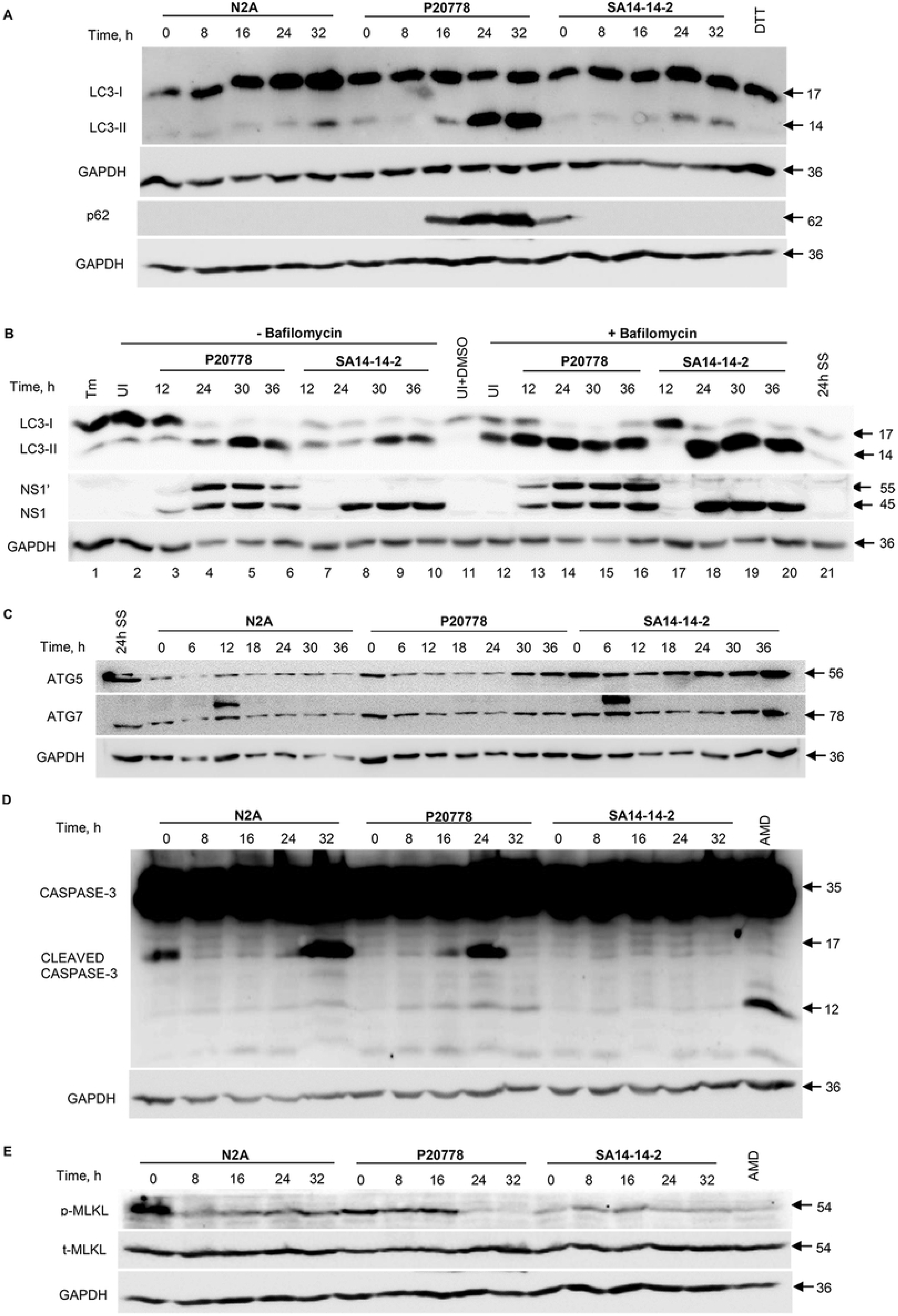
Differential modulation of autophagy by WT and vaccine strains of JEV. Lysates of N2A cells infected with P20778 or SA14-14-2 at a multiplicity of 10 were electrophoresed on SDS-10 % PAGE and transferred onto nitrocellulose membrane. Immunoblotting with antibodies specific to (A) LC3B, p62 and GAPDH, (B) LC3B, JEV NS1 and GAPDH after infected N2A cells were either left untreated or treated for 4 h with 100 nM bafilomycin A before being harvested at the indicated time points, (C) ATG5, ATG7 and GAPDH, (D) Caspase-3 and GAPDH and (E) pMLKL, tMLKL and GAPDH. UI-uninfected cell lysate; DTT-3 h treatment of cells with 10 mM DTT; Tm-treatment of cells for 5 h with 5 μg/ml tunicamycin; 24h SS-cells maintained for 24 h in serum free medium; AMD-cells treated for 18 h with 5 μg/ml AMD. Numbers on the right indicate sizes in kDa, of proteins detected in each panel.

We then investigated other death pathways in JEV-infected cells. While P20778 did not affect caspase-3 activation, we observed definitive suppression of residual caspase-3 cleavage in SA14-14-2-infected cells (Fig 3D, S2 FigF). P20778 infected cells revealed on the other hand, enhanced phosphorylation of MLKL, indicating early commitment to necroptosis following WT JEV infection (Fig 3E, S2 FigG). Both viruses also induced over expression of PARP-1 protein with minor and comparable enhancement of PARP-1 cleavage (S2 FigB).

### Differential stability of envelope proteins of WT and vaccine strains of JEV

The observed persistent eIF2α phosphorylation combined with IRE-1α activation and increased autophagy flux pointed to unresolved protein folding stress within SA14-14-2-infected cells. We therefore investigated levels of ER chaperones in virus infected cells. We observed dramatic upregulation of BiP and calnexin/calreticulin only in SA14-14-2-infected PS and N2A cells (Fig 4A and B). Keeping in mind the eight mutated residues of SA14-14-2 envelope, we proceeded to probe the stability and folding of envelope protein in cells infected with JEV.

**Figure 4.**
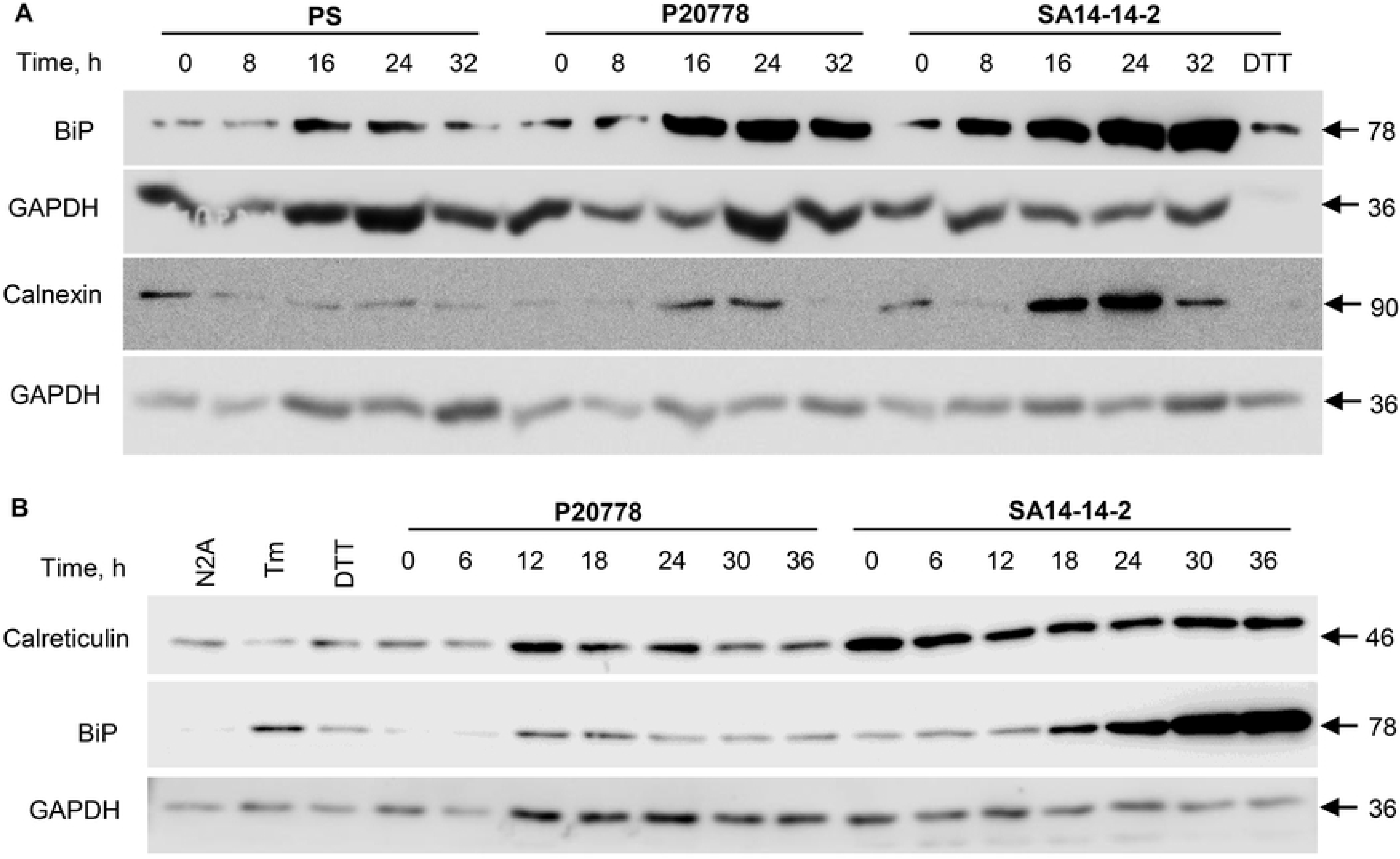
ER chaperone levels are increased in SA14-14-2 infected cells. Lysates of PS (A) and N2A (B) cells infected with P20778 or SA14-14-2 at a multiplicity of 10 and harvested at indicated time points were immunoblotted using antibodies specific to BiP, calnexin, calreticulin and GAPDH as indicated. N2A-uninfected N2A cell lysate; DTT-3 h treatment of cells with 10 mM DTT; Tm-treatment of cells for 5 h with 5 μg/ml tunicamycin. Numbers on the right indicate sizes in kDa, of proteins detected in each panel.

We utilized a rabbit polyclonal antiserum specific to JEV E protein to determine the half-life of E in cells infected with P20778 and SA14-14-2 strains of JEV. While WT E displayed a half-life of 5.9 h (S3 FigA, lanes 2 to 8; S3 FigB), SA14-14-2 E was rapidly degraded with half-life of 1.2 h pointing to its inherent instability (S3 FigA, lanes 9 to 15; S3 FigC). This was borne out by loss of conformational epitopes of SA14-14-2 E upon treatment with 7 M urea in contrast to P20778 E (S4 FigA). 7 M urea treatment did not affect reactivity of linear epitopes of SA14-14-2 E recognized by mAb CE3, F4B, DF4 and D10A, while that to conformational epitopes was lost (S4 FigA, lane 6) with the exception of 2D5. Reduction of proteins with DTT led to loss of all conformational epitopes in P20778 E protein also (S6 FigA, lane 3), with 1A5 and 2A9 being exceptions, indicating reliance on disulphide bond formation for generation of most of the conformational epitopes of E. A defect/delay in formation of disulphide bonds during the folding of SA14-14-2 E protein was evident from these observations. Following a short 5 min metabolic labeling of P20778-infected cells, use of N-ethyl maleimide to alkylate free sulfhydryl groups from the twelve cysteine residues of DTT-reduced nascent E protein to trap various folding intermediates revealed a fully reduced (R1) and two distinct intermediate forms (Ri and R2) along with a fully oxidized form (Ox) of E, the latter confirmed by H_2_O_2_ treatment (S4 FigB, lane 2). The rabbit polyclonal serum failed to immunoprecipitate the alkylated fully reduced form, presumably owing to occlusion of epitopes by alkylation (compare panels IP and Lysate in S4 FigB, lanes 3 to 8).

NEM alkylation of JEV-infected cell monolayers after 15 min of metabolic labeling revealed that in contrast to P20778 E, nascent SA14-14-2 E protein remained in various reduced forms for a much longer period post synthesis (S4 FigC). Each of the monoclonal antibodies mentioned above did indeed recognize its cognate epitope on the different reduced/oxidized forms of SA14-14-2 E protein (S4 FigC). The differential preference of the monoclonal antibodies for recognizing the different reduced and single oxidized form of SA14-14-2 E protein was also evident. More importantly, we also observed that several envelope-specific linear and conformational epitopes detected on extracellular virus particles of P20778 were missing on SA14-14-2 particles with the exception of the linear epitope recognized by D10A (S4 FigD).

### Belated folding kinetics of mutated SA14-14-2 envelope protein

We resorted to direct electrophoresis of labeled and alkylated infected cell lysates to follow the folding of E protein owing to poor ability to immunoprecipitated alkylated forms by the polyclonal anti-E serum (S4 FigB). Use of AMD prevented label incorporation into host proteins, allowing unambiguous visualization of the various forms of E (S4 FigB, lower panel). The E protein of JEV is co-translationally secreted into the lumen of the ER where it undergoes oxidation of its twelve cysteine sulfhydryl groups during folding. Wash out of DTT used to reduce metabolically labeled nascent E protein resulted in rapid oxidation (Ox) of cysteine sulfhydryl groups of fully reduced R1 form of P20778 E protein through an intermediate partially reduced species R2 (Fig 5A), leading to 45 % of the nascent protein being converted to the Ox form within 30 min of DTT removal (Fig 5A, lane 9). The precursor to product relationship of R1 and Ox forms of E through the intermediate R2 form was evident (Fig 5A, lanes 3 to 11; S6 FigB). In contrast, we saw a mere 19 % of label in the oxidized form of SA14-14-2 E protein at 30 min post DTT removal (Fig 5B, lane 9). Substantial proportion of labeled E protein remaining in the abundant fully reduced R1 form was in fact evidently degraded within this time, elevating the proportion of the scant quantity of Ox form of E. This dramatic instability of the reduced forms R1 and R2 of SA14-14-2 E was evident when band intensities were quantitated, with 80 % loss over a span of 60 min (S6 FigC), contributing to the observed short half-life of SA14-14-2 E protein (S3 Fig). It is to be noted that other viral proteins such as cytoplasmically localized NS3 and NS5 of SA14-14-2 were not similarly degraded (S5 FigB). Importantly, data in Fig 3B reveals that the ER luminally localized NS1 protein of SA14-14-2 is also spared from degradation observed for E, suggesting that the SA14-14-2 E was specifically targeted presumably by ER-associated degradation (ERAD) machinery of infected cells owing to its tardy folding kinetics. We observed no such differential instability of P20778 E relative to NS3 and NS5 (S5 FigA).

**Figure 5.**
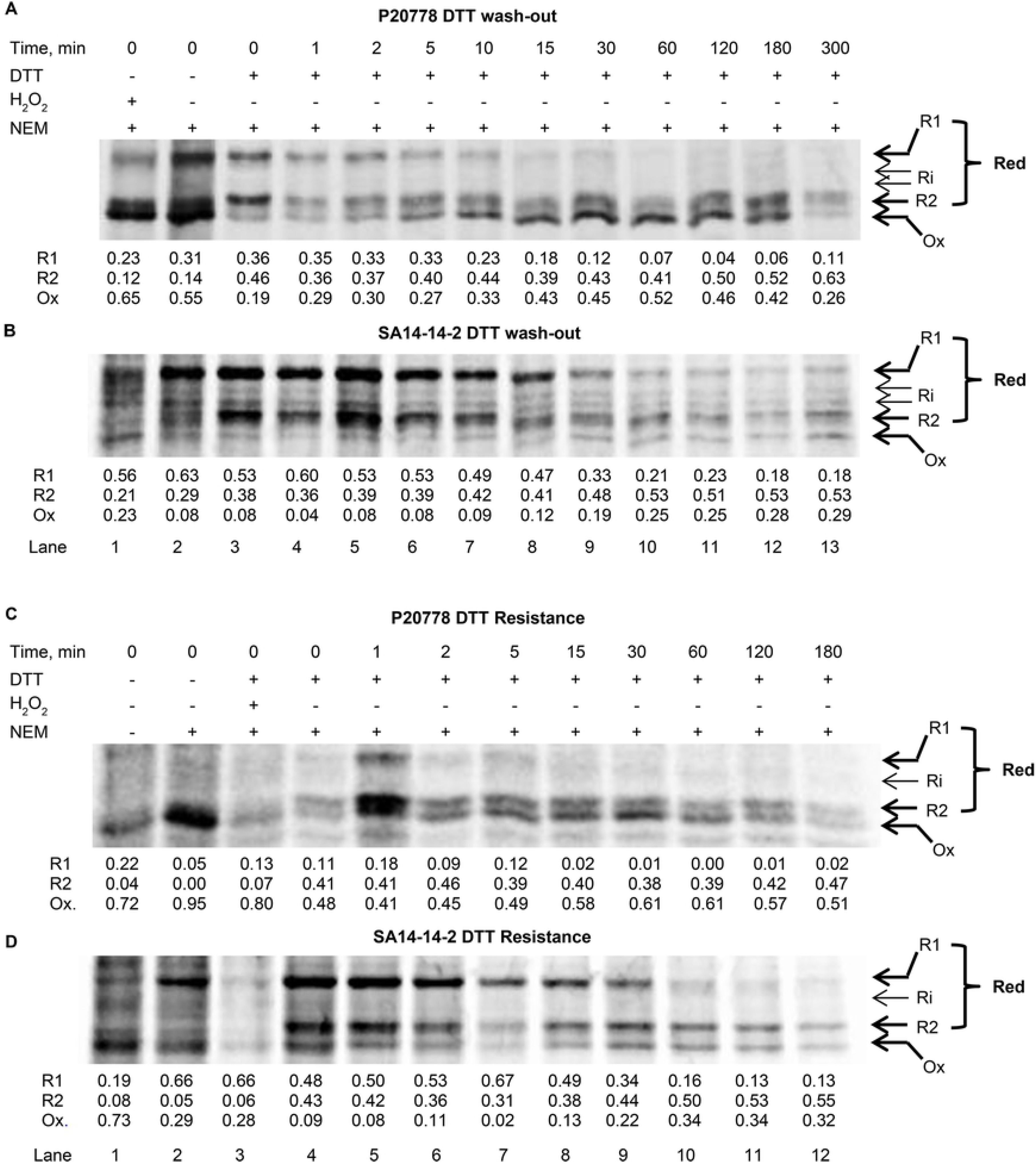
Folding kinetics of JEV E protein. PS cells infected with JEV P20778 (A) or SA14-14-2 (B) for 18 h were metabolically labeled for 5 min with ^35^ S-labeled methionine and cysteine followed by treatment with 10 mM DTT for 5 min to reduce cysteine sulphydryl groups on proteins. DTT was rapidly and extensively washed out and cells were reincubated in complete warm medium for the indicated times. Cell monolayers were treated with 20 mM NEM on ice for 10 min and lysates prepared without DTT were electrophoresed on SDS-7.5 % PAGE. PS cells infected with JEV P20778 (C) or SA14-14-2 (D) for 18 h were metabolically labeled for 5 min followed by replacement of medium with warm MEM containing 20 mM each of cysteine and methionine. Cell monolayers were treated with 100 mM DTT for 5 min at indicated time-points of chase, washed, alkylated with 20 mM NEM on ice for 10 min and lysates were electrophoresed on SDS-7.5 % PAGE. H_2_O_2_ was used at 100 μM for 5 min. The R1, Ri and R2 reduced along with oxidized form Ox of E are denoted by arrows. Numbers below the lanes provide the proportion of label in each form of E as denoted.

When we investigated the susceptibility of nascent E protein to reduction by DTT, we found rapid acquisition of resistance to DTT-mediated reduction by P20778 E within 15 min of synthesis (Fig 5C, lane 8; S6 FigD). In contrast, majority of SA14-14-2 E protein molecules remained susceptible to DTT reduction for at least an hour following its synthesis (Fig 5D, lanes 4 to 10) by which time, nearly 85 % of molecules vulnerable to reduction were degraded (Fig 5D, lane 10; S6 FigE). One min post synthesis, while a mere 18 % of WT E could be reduced to R1, this proportion was 50 % for SA14-14-2 E (Fig 5C and D, compare lanes 5). By 2 to 5 min post synthesis, nascent P20778 E was almost completely resistant to reduction to the fully reduced R1 form (Fig 5C, lanes 6,7). In contrast, a third of the residual degradation-resistant SA14-14-2 E protein could be reduced to R1 form even at 30 min post synthesis (Fig 5D, lane 9). Thus, mutations in the SA14-14-2 E protein rendered it incapable of achieving rapid oxidation-associated folding within the ER. Brefeldin treatment prior to metabolic labeling caused no difference in the kinetics of oxidative folding of WT and vaccine E proteins, testifying to ER as the site of E protein oxidation (data not shown).

### Accelerated furin cleavage of SA14-14-2 prM protein

Flavivirus maturation involves low pH-dependent furin mediated cleavage of the prM protein in the Golgi apparatus during egress [37, 38]. In extracellular virus particles of multiple flaviviruses, a sizeable proportion of prM has been reported to remain unprocessed by furin [37, 39, 40]. We observed approximately half the prM protein of extracellular P20778 virus particles had undergone furin cleavage (Fig 6A, fractions 1 to 4, lower panel). A small proportion of virus particles with maximum density recovered from the bottom of 70% sucrose following ultracentrifugation alone revealed complete processing of prM to pr and M (Fig 6A, fraction 6, lower panel). In contrast, we observed near total cleavage of prM to pr and M on extracellular SA14-14-2 virus particles with uncleaved prM barely visible even on long exposure (Fig 6A, lower panel). We surmised that the incomplete folding of the mutated envelope protein perhaps allowed enhanced access to the furin cleavage site of SA14-14-2 prM protein, which collectively led to the observed alteration of surface epitopes on virus particles of SA14-14-2 (S4 FigD). As expected, intracellular prM remained uncleaved in cell lysates of both viruses (Fig 6B, lower right panel)

**Figure 6.**
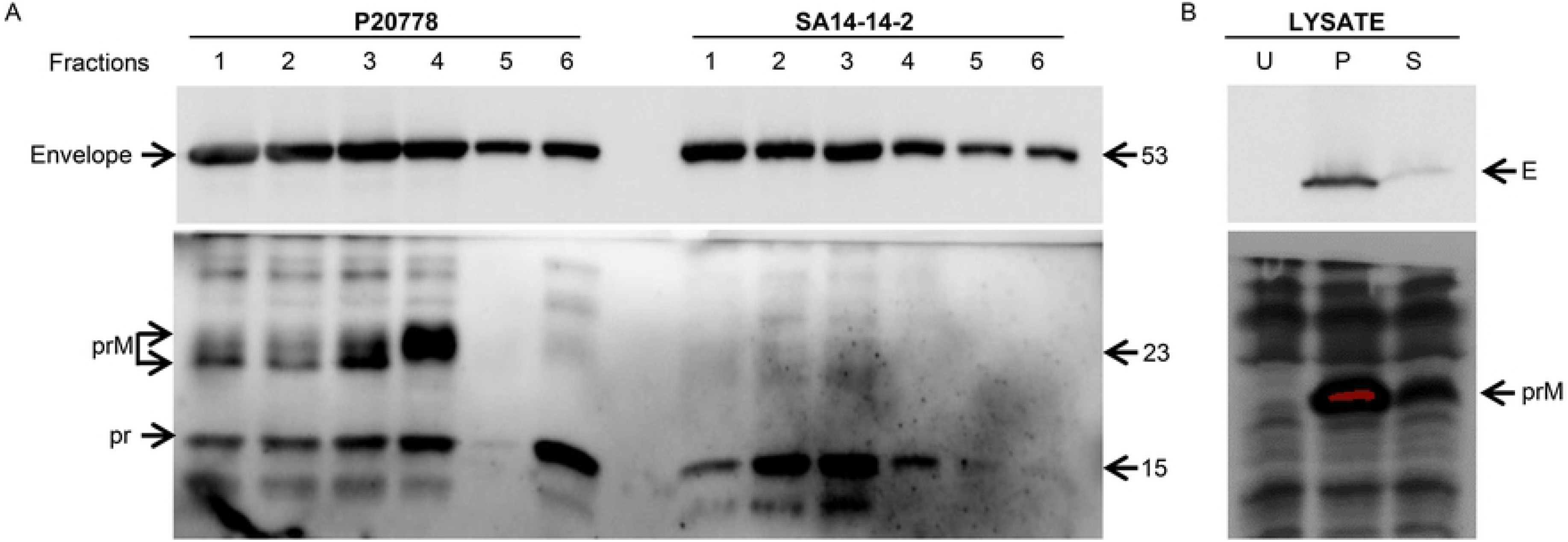
Efficient furin cleavage of SA14-14-2 prM protein. (A) Extracellular virus particles of P20778 and SA14-14-2 were ultracentrifuged on a 30 and 70 % discontinuous sucrose gradient as described in STAR Methods. Fractions from the interface of 30 and 70 % sucrose (lanes 1 to 4) and from the 70 % sucrose (lanes 5 and 6) were electrophored on SDS-12.5 % PAGE and immunoblotted with antibodies specific to the envelope (CE3, top panel) or prM (1C8, lower panel) proteins. The envelope, prM and cleaved pr proteins are indicated on the left with their sizes in kDa on the right. (B) Cell lysates of N2A left uninfected (U) or infected with P20778 (P) and SA14-14-2 (S) immunoblotted with antibodies specific to the envelope (CE3, top panel) or prM (1C8, lower panel).

### Enhanced CD8^+^ T cell responses in SA14-14-2 vaccinated individuals

In order to ask if the above differences in the cell biology of infection with P20778 and SA14-14-2 would be reflected in the immune response to the virus, we compared recall T cell responses in individuals exposed to circulating wild type JEV in endemic regions with those vaccinated with SA14-14-2. Capsid and E protein-specific T cells were both characterized by the dominance of CD4^+^ over CD8^+^ subsets in those naturally infected with circulating WT strains of JEV (HV and JEV groups; Fig 7A, top and middle panels). Capsid-specific CD4^+^ T cell responses were comparable between naturally infected healthy volunteers (HV) or recovered JEV patients (JEV) and vaccinated individuals (VAC; Fig 7A, top panel) while envelope specific CD4^+^ T cell responses were elevated in SA14-14-2 vaccinated individuals (VAC) relative to naturally infected individuals HV or JEV (Fig 7A, middle panel). Importantly, significantly greater percentages of capsid-specific CD8^+^ T cells secreting IFN-γ, TNF-α or MIP-1β including polyfunctional ones were observed in vaccinated individuals compared to infected HV and recovered JEV patients (Fig 7A, top panel). E-specific CD8^+^ T cells secreting IFN-γ, TNF-α and MIP-1β including polyfunctional T cells showed enhancement in SA14-14-2 vaccinated individuals relative to recovered JEV patients (Fig 7A, middle panel), in keeping with the earlier reported absence of JEV-specific CD8^+^ T cells in recovered JEV patients [41, 42]. In contrast, we did not observe such enhancement of NS3-specific T cells in vaccinees compared to naturally infected individuals (Fig 7A, lower panel). NS3 is known to be the strongest stimulator of human CD8^+^ T cells in JEV-endemic cohorts [42, 43]. IL-2 responses to JEV proteins were relatively weak (data not shown). These results suggested superior presentation of viral structural proteins to CD8^+^ T cells following SA14-14-2 infection.

**Figure 7.**
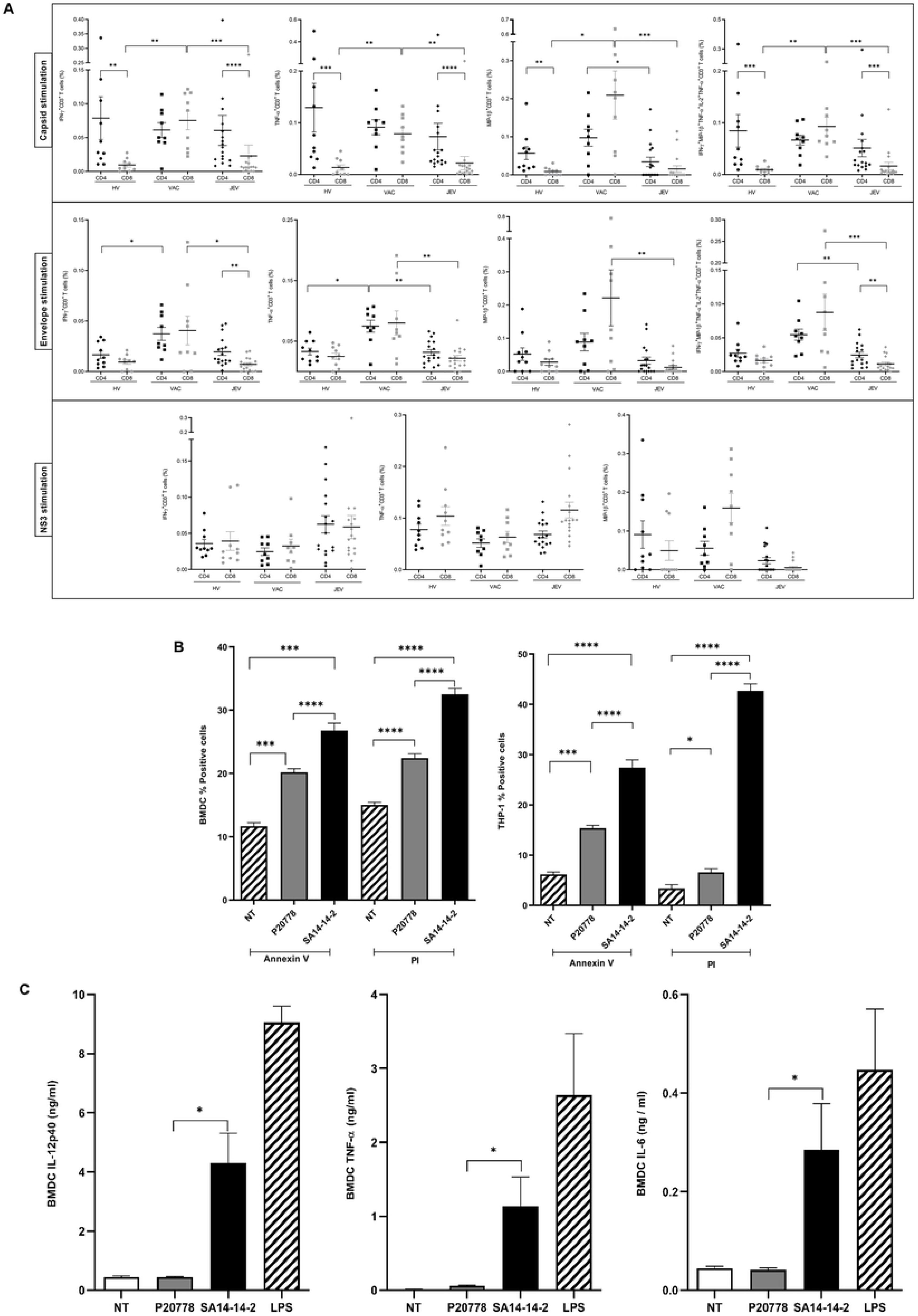
Superior immune activation by SA14-14-2. (A) Percentage of CD3^+^ CD4^+^ and CD3^+^ CD8^+^ T cells secreting IFN-γ, TNF-α or MIP-1β in response to stimulation with capsid (top panel), envelope (middle panel) and NS3 (lower panel) proteins of JEV P20778 compiled for healthy JEV infected volunteers (HV; N=10), SA14-14-2 vaccinated individuals (VAC; N=9) and recovered JEV patient (JEV; N=17) groups with median and IQR reported. Polyfunctional CD3^+^ CD4^+^ and CD3^+^ CD8^+^ T cells secreting a combination of IFN-γ, TNF-α, IL-2 and MIP-1β in response to capsid and envelope are also shown. Significance between CD4^+^ and CD8^+^ T cells within any group was determined using Mann Whitney U test while differences between CD4/CD8 T cells across the 3 groups were tested using Kruskal-Wallis H test with Dunn’s correction for multiple comparisons. (B) Enhanced cell death in SA14-14-2 infected antigen presenting cells. BMDC (left panel) and human THP-1 monocytes (right panel) were left uninfected or infected at 1 m.o.i. with P20778 or SA14-14-2 for 24 h prior to staining with Annexin V-FITC and propidium iodide (PI) as described in STAR Methods. Percentage positive cells are shown. N=4. Significant differences between groups were tested using ANOVA with Bonferroni correction for multiple comparisons. (C) Secretion of cytokines IL-12p40, TNF-α and IL-6 from BMDC infected with P20778 or SA14-14-2 for 12 h. NT-untreated; LPS-lipopolysaccharide from E. coli used at 0.1 μg/ml for 8 h. N=4. Significant difference between P20778 and SA14-14-2 strains of JEV was determined using Mann Whitney U test. P values are interpreted as follows: *, P ≤ 0.05; **, P ≤ 0.01; ***, P ≤ 0.001; ****, P ≤ 0.0001.

When we queried the ability of P20778 and SA14-14-2 to stimulate activation of antigen presenting cells, we observed enhanced death of primary mouse dendritic cells and human monocyte cell line THP-1 detected by Annexin-V and propidium iodide staining following 24 h infection by SA14-14-2 compared to P20778 (Fig 7B). In parallel, we also detected impressive increases in levels of inflammatory cytokines IL-12p40, IL-6 and TNF-α secreted from BMDC infected with SA14-14-2 compared to P20778 (Fig 7C). While we have not investigated the mode of death in infected cells, the above data suggest that presentation of viral antigens following infection with SA14-14-2 would be far more efficient relative to P20778. In light of the robust tropism of multiple mosquito-borne flaviviruses for human dendritic cells [44–46], we surmised that enhanced autophagy within antigen presenting cells infected with SA14-14-2 would engender superior cross presentation of viral antigens to CD8^+^ T cells as reported in the mouse model of influenza A virus infection [47].

## DISCUSSION

Both live attenuated vaccines available for two flaviviral pathogens, namely Yellow Fever 17D and JEV-SA14-14-2, were developed by serial passages in cultured cells, a process that led to accumulation of multiple mutations. Identity of and underlying molecular mechanisms triggered by the specific mutations responsible for their attenuated phenotype can help to exploit them for rationally developing vaccines against several other viral pathogens. We had the benefit of previous published studies that attributed the attenuation of SA14-14-2 to mutations in envelope and NS1’ [7–11]. Our previous studies had also revealed that the strongest correlate of naturally acquired immune protection in JEV-endemic human cohorts was flavivirus cross reactive CD8^+^ cytotoxic T cells secreting IFN-γ in addition to other TH1 cytokines such as IL-2 and TNF-α [41–43]. We therefore compared the cell biology of infections by WT and vaccine strains of JEV in cultured cells and attempted to relate them to differences in activation of antigen presenting cells as well as human recall T cell responses induced by the two strains of JEV. This rewarding exercise using simple approaches revealed the role of JEV NS1’ protein that is missing in SA14-14-2 in expropriating the host cell’s translational machinery to achieve efficient synthesis of viral proteins. NS1’ accomplished this by maintaining eIF2α in the dephosphorylated state both by upstream PERK destruction and downstream CHOP-mediated dephosphorylation. Increased levels of CHOP in JEV RP-9 infected BHK-21 and NT-2 cells was reported previously [48]. CHOP protein translation from its upstream ORF-containing mRNA requires the presence of phosphorylated eIF2α [29, 49]. The dramatically increased levels of CHOP in P20778-infected cells even in the absence of phosphorylated eIF2α is therefore surprising. Independent expression using lentiviral transduction confirmed NS1’ as the mediator of this phenomenon. The host proteins that NS1’ might interact with and the mechanism by which it upregulates CHOP expression await unraveling by future investigations. JEV NS4B was earlier reported to activate PERK by inducing its dimerization, leading to apoptosis and thereby encephalitis in mice [50]. That JEV targets PERK via multiple viral proteins in different tissues testifies to the importance of PERK in orchestrating both pro- and antiviral mechanisms in response to JEV infection. The reported reduction of JEV-induced apoptosis brought about by PERK inhibitor GSK2606414 in this study suggests that NS1’ may also serve to mitigate damage to neuronal cells caused by NS4B-induced PERK activation.

Most successful pathogens modulate death pathways in infected cells to escape host protective immune responses [51, 52]. Inhibition of autophagy to subvert host-protective immunity is a common strategy adopted by multiple pathogens through varied mechanisms [23]. WT JEV specifically prevented autophagy maturation leading to dramatic stabilization of LC3-II and p62 during late stages of infection while increasing autophagy during early phase of infection as also reported previously [31]. The stabilization of LC3-II at 48 but not 24 hrs post infection of N2A cells with JEV was also evident in Fig 3A of an earlier study [32]. The progressive loss of Beclin in P20778-infected PS cells suggests that JEV targets Beclin to prevent formation of autophagy maturation complex as reported for influenza A virus also [33]. Increased MLKL phosphorylation pointed to necroptosis in WT JEV infected cells. In light of the reported ability of the autophagosomal component p62 in recruiting RIPK1, leading to necrosome assembly in association with the autophagy machinery [53], necroptosis in WT JEV infected cells may be assisted by the observed p62 accumulation. In contrast, the enhanced autophagy in SA14-14-2 infected cells is most likely triggered by sluggish folding of E, with resultant unresolved UPR evidenced by high levels of ER chaperones and persistent eIF2α phosphorylation. These dramatic differences in death pathways activated by WT and vaccine strains of JEV would undoubtedly alter the host immune response to the two virus strains. Autophagy within antigen presenting cells has been reported to enhance cross presentation of viral antigens to prime CD8^+^ T cells [47]. Another study also proposed that autophagy-dependent antigen presentation on endocytosed cell surface MHC class I relies on TAP-independent vacuolar pathway where the acidic autophagolysosomal compartment would potentially stabilize MHC-peptide complexes [24, 54]. Since numerous viral immune evasion mechanisms target the TAP-dependent pathway of antigen presentation to escape CD8^+^ T cell-mediated antiviral immunity, autophagy dependent antigen presentation by MHC class I molecules allows circumventing this conventional pathway to achieve efficient priming of CD8^+^ T cells. The attenuating mutations in SA14-14-2 appear to have attained this outcome by disabling the WT virus’s mechanism(s) for inhibiting autophagy maturation. We therefore explored mechanisms instigated by mutations in the envelope protein of SA14-14-2 to enhance autophagy.

The retarded folding kinetics brought about by the numerous mutations in E of SA14-14-2 along with the observed accelerated furin cleavage of SA14-14-2 prM, culminated in dramatically altered surface epitopes on extracellular virus particles of SA14-14-2. We leveraged a panel of monoclonal antibodies that recognize several conformational and linear epitopes on JEV E along with alkylation of metabolically labeled nascent envelope protein to query the stability and folding kinetics of wild type and vaccine E proteins. The use of AMD to suppress host translation in virus-infected cells allowed us to effectively visualize and resolve the three different reduced forms along with oxidized form of the flaviviral E protein for the first time. The fully reduced R1 and oxidized forms of E revealed a clear precursor-product relationship during its folding, transiting through intermediate reduced (Ri) and a partially oxidized/reduced form R2. In addition to delay in folding of nascent SA14-14-2 E protein, we also observed continued vulnerability of its oxidized form to reduction by DTT for nearly an hour after synthesis; the P20778 E protein in contrast achieved DTT resistance in 5 min post synthesis. Failure of the mutated SA14-14-2 E protein to efficiently attain the stable oxidized form despite induction of ER chaperones BiP and calnexin/calreticulin probably resulted in its rapid degradation, most likely by the ER-associated degradation (ERAD) pathway. The acute stress inflicted on the ER protein folding machinery by mutated E of SA14-14-2 appears to be the immediate cause of autophagy enhancement mediated by persistent eIF2α phosphorylation. The abundant levels of viral NS proteins 1, 3 and 5 in SA14-14-2 infected cells (Fig 3B, S1 FigA) would suggest that despite IRE-1α activation, viral mRNA degradation by regulated IRE-1α dependent decay (RIDD) was not triggered. Among these NS proteins, it is particularly notable that NS1 of SA14-14-2 escapes degradation despite being ER localized (Fig 3B), further confirming that the targeted susceptibility of SA14-14-2 E to degradation emanates from its delayed folding kinetics within the ER.

We suspect that the unstable conformation of SA14-14-2 E protein permits superior access for golgi-resident furin to the prM protein in the immature viral particles on their exit path to the cell’s exterior. Extracellular virus particles of both JEV and WNV were shown to carry substantial proportions of uncleaved prM protein [39, 40]. Elegant cryo-electron microscopy studies combined with immunoprecipitation of metabolically labeled extracellular dengue virus particles with envelope and pr-specific antibodies revealed 30 to 40 % uncleaved prM distributed on 90 % of total virus particles, giving rise to a sizeable proportion of “partially mature” structurally dynamic virus particles [37]. The conformational flexibility and size heterogeneity of virus particles which was modulated by prM cleavage [55] would most likely permit the particles to “breathe” [56], rendering them difficult to target by the host immune system [57]. One is tempted to speculate that the near total cleavage of prM to pr and M on virus particles of SA14-14-2 that we observed, would likely give rise to homogeneous virus particles with rigid conformation that are efficiently targeted by host immune mechanisms. Furin cleavage of prM along with the acidic trans golgi environment are prerequisites for converting the intracellular immature ‘spiky’ virus particles into a flattened conformation of smooth mature particles [38, 58]. Structure investigations of SA14-14-2 virus particles ought to clarify whether the nearly complete prM cleavage renders these particles smooth and homogeneous in appearance.

The observed significantly higher levels of inflammatory cytokines from primary BMDC infected with SA14-14-2 as well as significantly greater death of infected BMDC and THP-1 human monocytes compared to P20778, most likely resulted from the enhanced autophagy triggered by SA14-14-2. This ability of SA14-14-2 to better activate BMDC was also evident when comparing two earlier reports [59, 60]. In keeping with this observation, we also noted enhanced CD8^+^ T cell responses directed to the structural proteins envelope and capsid of JEV in SA14-14-2 vaccinated individuals. The localization of capsid and envelope which are tethered to the ER membrane, would allow them access to the autophagy vesicles that are ER membrane derived [61, 62] and thus permit effective cross presentation to CD8^+^ T cells. This pathway would not be available for the cytoplasmically localized NS3, the dominant target of CD8^+^ T cells during natural infections, explaining the observed comparable NS3-specific T cell responses between naturally infected and vaccinated individuals. It is to be noted that the elevated CD8^+^ T cells were not evident in vaccinated individuals within the first 6 weeks post vaccination [63], suggesting efficient memory T cell generation following SA14-14-2 infection. It may be argued that these differences in CD8^+^ T cells between vaccinated and naturally infected individuals might merely reflect the time elapsed since infection. We cannot precisely determine the time of last exposure in naturally infected individuals residing in JEV-endemic regions. In fact, at the time of our sampling, JEV circulation was evident from the large number of hospitalized JE patients, suggesting that the lower levels of CD8^+^ T cells in naturally infected individuals was unlikely due to longer time elapsed between exposure to JEV and sampling. Our earlier studies revealed that during human infection with circulating strains of JEV, envelope protein predominantly stimulates CD4^+^ T cells with little evidence for CD8^+^ T cells [42]. WT JEV clearly possesses effective mechanisms to abort the generation of host-protective immune responses, including by suppressing the maturation of autophagic vesicles. Our results indicate that necroptosis induced by wild type JEV most likely suppressed presentation of CD8^+^ epitopes in the envelope and capsid proteins as also reported in HIV progressors [64], while the shift to autophagy induced by SA14-14-2 promoted cross presentation of these same epitopes to efficiently prime CD8^+^ T cells. Indeed, WT JEV has been reported to suppress priming of CD8^+^ T cells through the induced secretion of IL-10 by infected dendritic cells [44]. The use of recombinant envelope and capsid proteins derived from P20778 in our recall T cell assays testifies to the preservation of E-derived CD8^+^ epitopes in SA14-14-2 despite the multiple mutations. Thus, our findings throw light not only on mechanisms underlying the vaccine efficacy of SA14-14-2 but also illuminate the host immunity-subverting strategies adopted by WT JEV.

The development of existing live attenuated viral vaccines by serial passages of viral pathogens in cultured cells rendered it difficult to pinpoint those mutations that dictate the attenuated phenotype. The dominant attenuating effect of the envelope mutations revealed by substituting the envelope gene of wild type India78 strain of JEV with that from SA14-14-2 [7], suggests that defective folding of viral glycoproteins and the ensuing unresolved ER stress can sufficiently alter the cell biology of infection and antigen presentation to guarantee vaccine efficacy. The emphasis on JE virus neutralizing antibody titer for 50 % virus neutralization (PRNT50) of ≥ 10 as a sole surrogate of protection [65] led to a dearth of literature documenting T cell responses to JEV proteins in recipients of various JEV vaccines including SA14-14-2 [63]. Consequently, we know little about the extent to which vaccine-elicited CMI responses contribute to vaccine efficacy or reflect those seen in endemic settings. Interestingly, mice immunized with ChimeriVax-JE, in the complete absence of YFV-neutralizing antibodies, were protected against YFV challenge, confirming the autonomous protective ability of flavivirus-specific CMI responses [66] perhaps aided by non-neutralizing envelope specific antibodies. Conversely, the ability of neutralizing antibodies elicited by the mutated envelope of SA14-14-2 to effectively neutralize circulating wild type strains of JEV deserves scrutiny.

## MATERIALS AND METHODS

### Cell lines and viruses

Mouse neuroblast cell line Neuro-2a (CLS # 400394/p451_Neuro-2A, RRID:CVCL_0470) *Aedes albopictus* cell line, C6/36 (ATCC # CRL-1660, RRID:CVCL_Z230) and porcine kidney fibroblast cell line PS [67], obtained from the National Centre for Cell Science, Pune, India were grown in minimum essential medium (MEM; Gibco #41500-018) supplemented with 5% fetal bovine serum (FBS; Gibco #11573397). THP-1 human monocyte cells from American Type Culture Collection (ATCC) were maintained in RPMI 1640 medium (Gibco #11500456) with 10% heat inactivated FBS. Bone marrow derived dendritic cells were obtained by differentiating bone marrow cells from 6 week old BALB/c mice as described [68] and infected with JEV strains at a multiplicity of 1.

JEV strain P20778, West Nile Virus strain E101 (National Institute of Virology, Pune, India) and SA14-14-2 (Chengdu Institute of Biological Products, Chengdu, Sichuan, China) were propagated in the *Aedes albopictus* cell line, C6/36 or Neuro-2a cells infected at a multiplicity of infection (m.o.i) of 0.02. Virus stocks were harvested from the former after 6 days; P20778 and SA14-14-2 were harvested from the latter 72 and 96 hours post infection (h.p.i), respectively. Virus titres were determined by plaque assay on PS cells infected with serial dilutions of virus stocks; monolayers were stained with 1 % crystal violet in 20 % ethanol-water 72 to 96 h.p.i.

### Ethics Statement

This study was performed in accordance with the principles of the declaration of Helsinki. The study was approved by the IISc Institutional Human Ethics Committee (ref 5/2011). Vaccination of healthy volunteers with SA14-14-2 was registered at clinicaltrials.gov (https://clinicaltrials.gov/ct2/show/NCT01656200).

### Participants

Recovered JEV patients (JEV; N=17) were recruited at dedicated outpatient clinics held at the Vijayanagar Institute of Medical Sciences, Bellary, Karnataka while healthy donors (HV; N=10) were drawn from family members of patients and members of the local community as previously reported [42]. Healthy adults recruited by word of mouth and advertisement in Bangalore and vaccinated with SA14-14-2 (VAC; N=9) as reported earlier [63] were also used for this study. Patients and vaccinees were bled once 10 to 12 months post discharge/vaccination.

### Infection of cells and Lysate preparation

PS and N2A cells were infected with JEV strains at a multiplicity of 10 for all experiments. Pre-warmed growth medium was changed completely every 8 h. Cells were harvested by scraping the monolayer. Cell pellets were lysed in ice cold lysis buffer (20 mM Tris- HCl pH 7.5, 50 mM sodium pyrophosphate, 50 mM NaF, 150 mM NaCl, 1 mM EDTA, 1 mM EGTA, 100 μM Na_3_ VO_4_, 1 % Triton X-100) for 15 min. Lysates were clarified by centrifugation at 1000 rpm, 4 °C for 10 min and stored at −80 °C.

### Western blot analysis

Lysates prepared as mentioned above were electrophoresed on SDS-PAGE, transferred to nitrocellulose or PVDF membranes and western blotting was conducted using the following primary antibodies: eIF2α (Cell Signaling Technology #9722), p-eIF2α (Cell Signaling Technology #9721), ATG7 (Cell Signaling Technology #2631), ATG5 (Cell Signaling Technology #8540), LC3B (Santa cruz #sc-271625), β-actin (Cell Signaling Technology #4967), GAPDH (Santa cruz #sc-47724), CHOP (Cell Signaling Technology #2895S), Calnexin (Cell Signaling Technology #2679S), PERK (Cell Signaling Technology #3192), BiP (Cell Signaling Technology #3177), Calreticulin (Rabbit mAb D3E6; Cell Signaling Technology #12238), PARP-1 (Rabbit mAb 46D11; Cell Signaling Technology #9532), Caspase-3 (Rabbit mAb 8G10; Cell Signaling Technology #9665), ATF4 (Rabbit mAb D4B8; Cell Signaling Technology #11815) MLKL (Rabbit mAb D2I6N; Cell Signaling Technology #14993), pMLKL (phospho S345; Abcam #ab196436), Beclin-1 (Cell Signaling Technology #3738), IRE-1α (Novus Biologicals #NB100-2323), XBP-1 (Rabbit Polyclonal; Novus Biologicals #NBP1-77681), SQSTM1/p62 (D5E2; Cell Signaling Technology #8025 and rabbit polyclonal; Abcam #ab91526, to detect porcine and murine p62, respectively). Antibodies to viral proteins E, prM, NS1’ (mouse monoclonal), NS3 (rabbit polyclonal) and NS5 (mouse polyclonal) were generated in house [43, 69]. Membranes blocked for 2 h at room temperature with 0.5% non-fat milk powder (Carnation) in TBS buffer (10 mM Tris pH-8, 150 mM NaCl) were incubated with primary antibody in TBS overnight at ambient temperature. Membranes were washed thrice with TBS/ 0.1% Tween20, incubated with appropriate secondary antibody in TBS buffer, washed thrice with TBS/ 0.1% Tween20, and developed using BIO-RAD Clarity western ECL substrate (#170-5060) using ImageQuant™ LAS 4000 from GE healthcare Life Sciences.

### Cloning of NS1’ and NS1 in lentiviral vector

Total RNA obtained from JEV P20778-infected PS cells harvested 24 h.p.i. was reverse transcribed using primer OSV 381 with AMV Reverse Transcriptase (Promega). The JEV NS1 gene was PCR-amplified with Deep Vent polymerase (New England Biolabs) using forward primer OSV 389 and sequential reverse primers OSV 381, OSV 382, OSV 387, OSV 388, OSV 390 and OSV 391 (Table S1) to obtain the full length NS1’ gene. OSV 387 and OSV388 were designed to disrupt the slippery heptanucleotide and pseudoknot structure at the beginning of the NS2a gene of JEV that promote ribosomal frameshifting, without altering the amino acid sequence of the NS1’ C-terminus. The full length NS1’ gene was amplified using forward primer OSV 389 and reverse primer OSV393 containing the haemagglutinin (HA) tag sequence followed by a termination codon and a *Not* I site. Cohesive ends were generated by restriction digestion of this PCR product using *Eco* R1 and *Not*1 and ligated using T4 DNA ligase (Promega) with *Eco* R1, *Not*1 digested and Calf Intestinal Phosphatase treated pCDH-CMV-MCS-EF1α-copGFP Dual Promoter Cloning and Expression Lentivector (System Biosciences LLC, #CD511B-1). Similarly, cDNA synthesized using reverse primer OSV278 was used to amplify the NS1 gene along with forward primer OSV389 (Table S1). The PCR product was digested with *Sal* I, Klenow filled and digested with *Eco*RI. Ligation to pCDH-CMV-MCS-EF1α-copGFP Dual Promoter Cloning and Expression Lentivector digested with *Not* I, Klenow filled and *Eco*RI digested was carried out. Recombinant plasmids from transformed *E. coli* DH10B electrocompetent cells identified by diagnostic restriction digestion were verified by sequencing. Expression of the authentic NS1’ and NS1 proteins in HEK-293T cells transfected with the recombinant pCDH-NS1’HA and pCDH-NS1 plasmids was confirmed by western blotting using a monoclonal antibody generated against a peptide sequence derived from the frameshifted C-terminal sequence of NS1’ (see below) which specifically detected NS1’ in P20778-infected cells and a polyclonal serum raised to *E. coli* expressed recombinant JEV-NS1 protein [43], respectively.

### NS1 and NS1’ Lentivirus Generation

293T/17 cells were transfected with pCDH-NS1’HA or pCDH-NS1 along with the three packaging plasmids psPAX, pVSV-G and pRSV-rev using calcium phosphate. Briefly, 0.6 million 293T/17 cells were seeded on 35 mm dish (BD Falcon) in 2 ml MEM, 5 % FBS. One day later, medium was changed one hour prior to transfection and fresh 1.8 ml complete MEM was added. Transfection mix containing the four plasmids in 50 μl autoclaved water, 50 μl 2.5 M calcium chloride and 100 μl 2X HEPES buffered saline was immediately added onto the 293T/17 cells drop by drop. Brief centrifugation for 120 s at 1000 rpm in a swing out rotor was carried out to enable rapid sedimentation of precipitates. 4 h post transfection, medium was changed with pre-warmed complete MEM. Supernatant collected 4 days post transfection was stored at −80 °C. p24 ELISA was performed to determine the lentivirus titer using Perkin Elmer p24 ELISA kit (#NEK050001KT) according to manufacturer’s instructions.

### Monoclonal antibody generation

To generate NS1’-specific monoclonal antibody a peptide sequence was derived from the C-terminal frameshifted segment of NS1’ protein (SQEVDGQIDHSCGFG) using Bcepred software (http://crdd.osdd.net/raghava/bcepred/) based on hydrophilicity, flexibility, accessibility, exposed surface, polarity and antigenic propensity. The peptide was conjugated to BSA using glutaraldehyde and used to immunize BALB/c mice (50 μg each per mouse, 3 times at 4-week intervals). To generate antibodies against JEV/WNV envelope/premembrane proteins, BALB/c mice were injected intraperitoneally with 2 × 10^6^ pfu of virus. Subsequently, two boosters with 2 × 10^6^ pfu of virus were administered at intervals of 30 days. 7 days post injection of each booster, sera were collected by retro-orbital bleeding and tested by western blotting of JEV-infected cell lysates and ELISA against the peptide with dilutions ranging from 1:1000 to 1:100,000. Spleen cells isolated from the mouse with the best serum titre were fused with Sp2/0 cells in the ratio of 5:1 using polyethylene glycol (PEG) 3000 (#817019, Merck). 10 million cells of the fusion mix were combined with 2x 10^4^ BALB/c peritoneal macrophages and seeded in a 96-well plate. Hybridoma were selected in HAT medium for 6 days followed by HT supplemented medium [70]. Culture supernatants were screened by ELISA on day 10. Culture supernatants collected from the monoclonal antibody producing cloned cells were verified by reactivity to authentic NS1’/E/prM in JEV-infected cell lysates by western blotting.

### Metabolic labelling and immunoprecipitation

Cells were seeded at a density of 2 × 10^5^ cells/35 mm dish and cultured for 48 h at 37 °C to reach 70 % confluence. Cells were infected at a multiplicity of 10 with P20778 or SA14-14-2 at 37 °C for 1 h. Virus inoculum was replaced with pre-warmed complete MEM. 15 h.p.i. for P20778 and 19 h.p.i. for SA14-14-2 endogenous pools of cysteine and methionine were depleted by treating cells for 2 h with Cys^−^ Met^−^ MEM (MP Biomedicals #1641454) containing 7.5 μg/ml Actinomycin-D (AMD; Sigma-Aldrich #A9415) to inhibit host transcription. Cells were pulsed with 450 μCi/35 mm dish of ^35^ S-labeled methionine and cysteine (American Radiolabeled Chemicals, 1175 Ci/mmol; #ARS0110A) diluted in 2 ml MEM, 1 % FBS and 7.5 μg/ml AMD for 5 min. For chase, radiolabel was removed and pre-warmed complete MEM containing cold 20 mM each of cysteine and methionine was added. Cells were either left untreated or treated with 100 mM dithiothreitol (DTT) for 5 min, 100 μM hydrogen peroxide (H_2_O_2_) for 5 min or alkylated with 20 mM N-ethyl maleimide (NEM; Sigma #E3876) in 100 mM sodium phosphate, 150 mM NaCl pH-7.2 (PBS) on ice for 10 min. Monolayers were harvested by scraping and cell pellets lysed either in lysis buffer (20 mM Tris pH-7.5, 150 mM NaCl, 1 mM EDTA, 1 mM EGTA, 1 % Triton X-100, 2.5 mM sodium pyrophosphate, 1 mM beta-glycerophosphate, 1 mM sodium vanadate, 1 mM sodium fluoride, 1X protease inhibitor cocktail) for electrophoresis on 7.5 % PAGE or in ice-cold RIPA lysis buffer (150 mM NaCl, 1 % Triton X-100, 0.5 % deoxycholic acid) for immunoprecipitation with Protein A/G beads coated with rabbit polyclonal/mouse monoclonal antibody against envelope protein. Bound proteins eluted from washed beads using 1X Laemmli SDS-PAGE buffer without DTT at 37°C for 10 min were electrophoresed in SDS-7.5 % PAGE. Gels were dried in a BIO-RAD Model 583 gel dryer and developed using the Typhoon FLA 9500 biomolecular imager (GE healthcare Life Sciences).

### ELISA

Peptides were coated at a concentration of 1 μg/ml, 100 μl/ well. 100 μl containing 10^4^ pfu of either P20778 or SA14-14-2 virus in 100 mM carbonate buffer pH-9.6 was coated on high binding ELISA plates (Orange scientific) and incubated for 12 h at 4 °C in a humidified chamber. Wells were washed thrice with 100 mM sodium phosphate, 150 mM NaCl, 0.05 % Tween 20 pH-7.2 (PBST), blocked with 200 μl 0.1 % BSA in 1X PBS and incubated at room temperature for 2 h. Wells were washed thrice with PBST followed by incubation with 100 μl of appropriate hybridoma culture supernatant at ambient temperature for 3 h. 100 μl of horseradish peroxidase (HRP)-conjugated goat anti-mouse secondary antibody (Southern Biotech #1036-05) was then added at a dilution of 1:5000 and incubated for 1 h at ambient temperature. Wells were washed thrice with PBST, once with PBS and 100 μl 1X TMB/H_2_O_2_ substrate was added. Reaction was stopped after 30 min using 0.2 N sulphuric acid and absorbance at 450 nm was obtained using a TECAN Infinite F50 ELISA reader with Magellan V 7.2 software. Mouse IL-12p40, IL-6 and TNF-α were measured using DuoSet® ELISA kits (R&D Systems).

### Intracellular cytokine detection

Whole, heparinized (sodium heparin) blood was diluted 1:1 with RPMI 1640 and 1 ml aliquots were stimulated with recombinant NS3, envelope or capsid proteins of JEV P20778 purified as reported [43] at a concentration of 5 μg/ml for 20 h as described [71]. Brefeldin A (Sigma-Aldrich #B7651; 10 μg/ml) was added 2 h after peptide addition while monensin (Sigma-Aldrich #475895) was added 6 h later at a concentration of 0.75 μM, a concentration that we determined to effectively block secretion of cytokines without adversely affecting cell viability following 12 h exposure. We used both these secretion inhibitors since they differentially block surface expression/secretion of several markers studied [72, 73]. 12 h later, erythrocytes were lysed by addition of 10 volumes of ammonium chloride lysis solution (166 mM ammonium chloride, 9.9 mM potassium bicarbonate and 0.126 mM EDTA), vigorously vortexed for 1 min and leukocytes retrieved by centrifugation were fixed using 2 % paraformaldehyde (Merck #158127) on ice for 10 min and washed with PBS-0.1 % sodium azide solution.

Cells were then permeabilized with 0.1 % saponin (Merck #47036) and 0.1 % bovine serum albumin in PBS for 15 min on ice and intracellular cytokines were detected using an antibody cocktail consisting of titrated amounts of anti-CD3-APC-H7 (SK7), anti-CD8-PerCP (SK1), anti-IFN-γ-PECy7 (B27), anti-IL-2-FITC (MQ1-17H12), anti-TNFα-APC (6401-1111) and anti-MIP-1β-PE (D21-1351), from BD Pharmingen, San Diego, CA. Data were acquired on a BD-FACS Canto II flow cytometer (Becton Dickinson, San Jose, CA). Singlet small lymphocytes were collected after excluding dead cells and debris by gating on forward versus side scatter and then gated on CD3^+^ T lymphocytes. CD3^+^ cells negative for CD8 were considered as CD4^+^ T cells (Figure S7). For each analysis, a minimum of 100,000 CD4^+^/CD8^+^ T cell subsets were acquired and data analyzed using FlowJo (Version 7.0 for Windows, Ashland, Oregon), PESTLE and SPICE [74] software. Antibody-stained unstimulated cells served as control. A positive response was defined by a minimum number of 50 events over controls. Gates were positioned to retain the response of unstimulated cells ≤ 0.01 % of total CD4^+^/CD8^+^ T cells for IFN-γ, TNF-α and IL-2 secreting T cells, while it was ≤ 0.05 % for MIP-1β single cytokine secreting T cells.

### Monitoring cell death by flow cytometry

Cells were stained with a combination of Annexin V-FITC (BD Biosciences #556420) and propidium iodide (BD Biosciences #556463) as described [75]. Cells washed in ice cold PBS were suspended in binding buffer (10 mM HEPES pH-7.4, 140 mM NaCl, 2.5 mM CaCl_2_) and stained with a combination of Annexin-V FITC and 250 ng of propidium iodide in 100 μl for 30 minutes. Cells were washed in binding buffer, fixed with 2 % paraformaldehyde (Merck #158127) for 10 min on ice, washed with cold PBS and treated with RNase A (0.1 mg/mL; ThermoFisher Scientific #EN0531) for 15 min at 37 °C before being acquired in a BD-FACS Canto flow cytometer.

### Sucrose gradient centrifugation of JEV

P20778 or SA14-14-2 virus grown on C6/36 cells were overlayed on a cushion of 8 ml 25 % sucrose and ultracentrifuged at 4 °C for 3 h at 80,000 × g in a Beckman Model L8-70M ultracentrifuge. Virus pellets resuspended in GTNE (200 mM glycine, 50 mM Tris pH 7.5, 100 mM NaCl, 1 mM EDTA) buffer were overlayed on a sucrose step gradient consisting of 1.5 ml each of 70 and 30 % sucrose and centrifuged at 4 °C for 3 h at 100,000 × g in a SW60 Ti rotor in a Beckman Model L8-70M ultracentrifuge. Virus band was observed at the 30-70 % sucrose interface. 0.5 ml fractions were collected from the 30-70 % sucrose interface until the bottom of the tube and stored at −80 °C.

### Statistical analysis

All western blotting experiments were carried out in biological triplicates. Statistical analyses were done using GraphPad Prism version 8.0. Significant difference between two or multiple groups was tested using Mann–Whitney *U* test (two-tailed) and non-parametric Kruskal-Wallis test with Dunn’s test or parametric ANOVA with Bonferroni correction for multiple comparisons, respectively. Band intensities quantitated using ImageJ were compared using unpaired Student’s *t* test with alpha set at 0.05.

## Acknowledgements

We thank Sai Pallavi Pradeep and Madhusudan Thirumallesh for help with compiling figures. We thank Sukanya Raghu for help with Western blots. This study was funded by a grant (EMR/2016/000373) awarded to VS by the Science and Engineering Research Board, Department of Science and Technology, Government of India and DBT-IISc Partnership Program Phase-II (BT/PR27952/INF/22/212/2018) awarded to the Biology Division, Indian Institute of Science. This research was also funded in part, by a Wellcome Trust fellowship awarded to LT [087757/Z/08/Z], currently funded by Wellcome trust grant number 205228/Z/16/Z. For the purpose of Open Access, the author (LT) has applied a CC BY public copyright licence to any Author Accepted Manuscript version arising from this submission. LT is also supported by the National Institute for Health Research Health Protection Research Unit (HPRU) in Emerging and Zoonotic Infections (NIHR200907) at University of Liverpool in partnership with Public Health England (PHE), in collaboration with Liverpool School of Tropical Medicine and the University of Oxford. LT is based at University of Liverpool. The views expressed are those of the author(s) and not necessarily those of the NHS, the NIHR, the Department of Health or Public Health England. The funders had no role in study design, data collection and analysis, decision to publish, or preparation of the manuscript.

## Author Contributions

VS conceived, designed, planned and executed experiments, supervised the study, curated, interpreted and visualized data, acquired funds and resources, wrote and edited the manuscript; LT conceived designed and executed human vaccination studies and acquired funds and resources; PHV executed experiments, interpreted data, prepared figures and wrote the manuscript; NK carried out and interpreted experiments, prepared figures; SK and KC carried out experiments.

## Data availability

All relevant data are within the manuscript and its Supporting Information files. De-identified flow cytometry files (.fcs) that support the results reported in this article, have been deposited in Mendeley Data (https://data.mendeley.com/datasets/2g9jv6bzx5/1).

